# Portable low-cost optical density meter

**DOI:** 10.1101/2021.05.14.444207

**Authors:** Robert R. Puccinelli, Joana P. Cabrera, Emily Huynh, Paul M. Lebel, Rafael Gómez-Sjöberg

## Abstract

Measuring optical density (OD) is a very common technique in biological laboratories to determine the concentration of a substance in solution or of bacteria (or microscopic particles) in suspension. For example, bacterial cultures engineered to produce (express) a protein or compound of interest are a workhorse of modern molecular biology laboratories. Commonly, the expression of the product is triggered (induced) by a chemical signal added to the culture at the proper time in the growth curve of the culture (typically in the middle of the exponential growth phase, at an OD value of ∼0.6). The most common tool for measuring OD is a spectrophotometer. However, most spectrophotometers are sophisticated, non-portable and expensive laboratory instruments, costing tens of thousands of dollars. Even a very low cost spectrophotometer for educational use costs at least US$1,000. Because of the cost, even well resourced labs have only one instrument, which becomes a bottleneck when multiple bacterial cultures need to be monitored simultaneously. The problem is more acute in developing countries, where multiple labs have to share a single spectrophotometer, or there’s no such instrument at all. Having a cheap and simple device to measure OD would enable multiple people in a laboratory to monitor their bacterial cultures independently, even in resource-limited settings. At the same time, a portable OD meter could be useful for field work. Here we present the detailed build instructions and characterization of a very simple OD meter that costs only US$60, and can measure OD values from ∼0.05 to 2.0.

**Specifications table:** 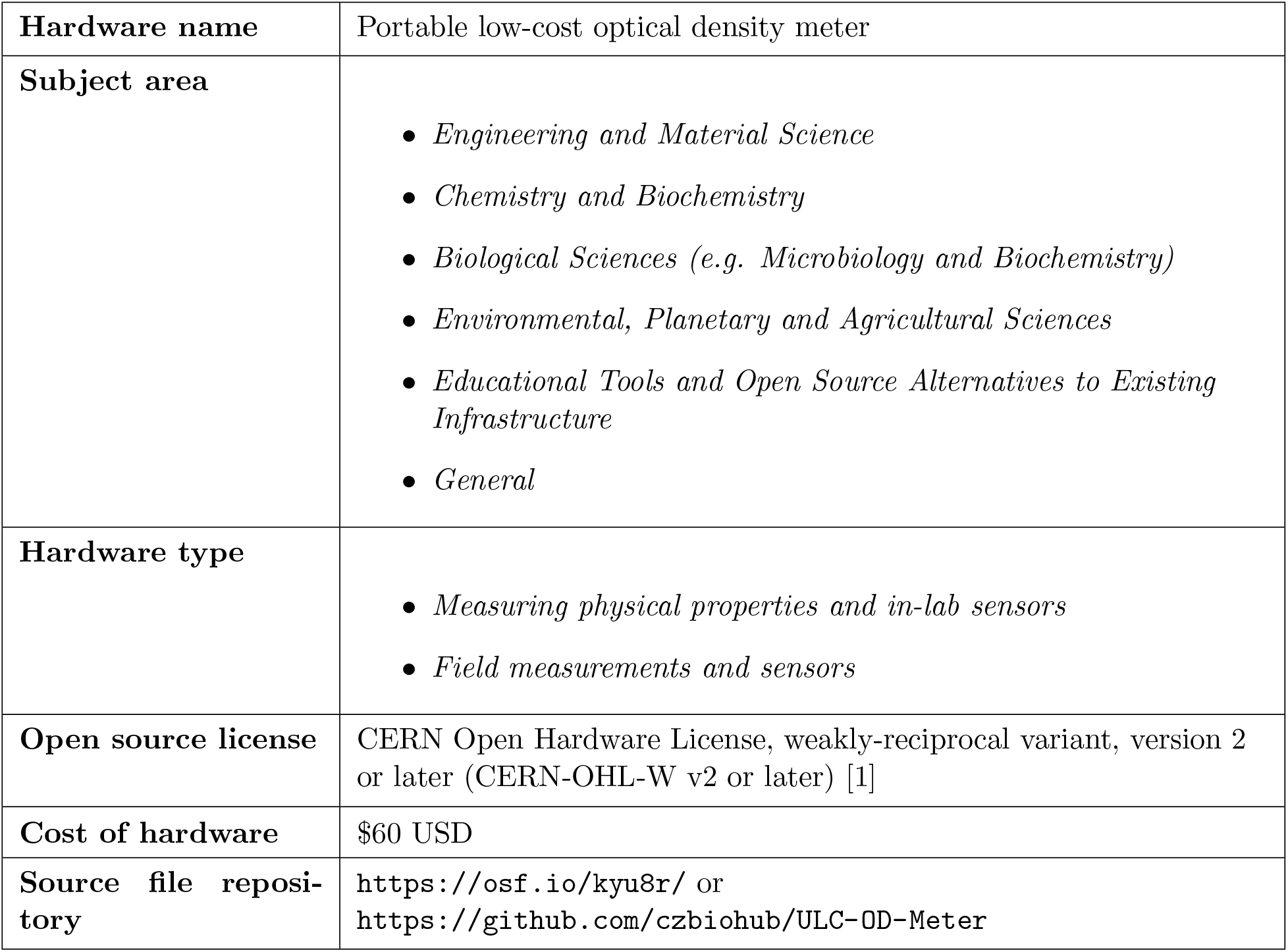

## 1. Hardware in context

Bacterial cultures engineered to produce (express) a protein or compound of interest are a workhorse of modern molecular biology laboratories. Commonly, the expression of the product is triggered (induced) by a chemical signal added to the culture at the proper time in the growth curve of the culture, typically in the middle of the exponential growth phase, at an OD value of ∼0.6. In order to monitor the growth of the culture, its OD is measured over time using a spectrophotometer, which is a very common tool in most laboratories, especially in industrialized countries. However, most spectrophotometers are sophisticated, non-portable and expensive laboratory instruments, costing tens of thousands of dollars. Because of the cost, even well resourced labs have only one instrument, which becomes a bottleneck when multiple cultures need to be monitored simultaneously, and laboratories in low-resource areas might not have a spectrophotometer at all. Even a very low cost spectrophotometer for educational use costs at least US$1,000, and a recently-published low-cost spectrophotometer costs ∼US$262[5].

In 2019, a researcher from the Chan Zuckerberg Biohub was in Kampala, Uganda, teaching a two-week long laboratory course, which needed the expression of particular proteins in engineered bacterial cultures. In the middle of the course, it was discovered that the spectrophotometer in the laboratory was broken and the induction on the cultures was being triggered too late in the growth curve. Repairing such a sophisticated instrument in Uganda can take several months, and without it the training course would be severely impaired. The trainer in Kampala contacted the Bioengineering team at the Biohub and asked if we could build a very simple device that would allow them to monitor the bacterial culture growth and salvage the course. In three days we designed and built a very low cost (US$60) portable optical density meter with a measurement wavelength of 590 nm and a validated operational range of 0.05 to ∼1 OD, which was carried to Uganda by another Biohub researcher that was already scheduled to fly there. Subsequently, the design was improved slightly and validated to extend the measurement range to 2.0 OD. While the device was originally designed to measure the growth curve of bacterial cultures, it can also be useful in other applications, such as determining the concentration of small particle suspensions, monitoring reactions that change the color of a solution, measuring the concentration of solutions of compounds that absorb strongly at 590 nm, or performing simple checks on the quality/turbidity of water.

## 2. Hardware description

The low-cost OD meter is compact and portable, measuring 114 mm long, 85 mm wide, and 42 mm high. It can be powered from any USB port, including those present in a standard cell phone charger or a rechargeable power pack, using a USB to micro-USB cable. It has a recess for a standard spectrophotometry cuvette (with a cap to keep ambient light out) and a high contrast LCD screen that displays measurements and user prompts. The device can measure OD on a range from ∼0.05 to 2.0 by measuring dark and blank reference samples to set the proper OD scale, and it also reports the raw values of the light intensity being measured. The dark and blank reference calibrations are very easy to do when measuring OD up to 1.0 (a 10x attenuation of the light) with solutions (non-scattering samples), and up to 2.0 (100x attenuation) with bacterial suspensions (scattering samples). To accurately measure solutions with OD values between 1.0 and 2.0 a more complex calibration technique is required, as discussed in the operation instructions section. The design is based on a low-cost Arduino microcontroller, whose programming can be easily changed by knowledgeable users to expand the capabilities of the meter (such as automatically sending the measurements to an external logging device). The LED used as a light source has a peak wavelength of 590 nm (yellow), with a full-width at half-maximum emission of ∼15 nm. This LED is aimed at the measuring region of the cuvette through a pupil that is 1.3 mm in diameter and 6 mm long, to restrict the angular spread of the emission. The LED light transmitted through the sample in the cuvette is measured by a photodiode located on the opposite side of the cuvette, facing the LED through a 1.3 mm diameter, 7 mm long pupil that is co-linear with the LED pupil.

This device joins a series of related low-cost instruments that have been reported in the literature for performing colorimetry [2, 7], measuring turbidity [4, 6], measuring the OD of radiographic film [3], and spectrophotometry [5]. Unlike some of these devices, which need an Android smart phone to operate, our OD meter is completely stand-alone as it has its own user interface and does not require any other device for operation, except for an external power supply, such as a battery pack.

This device is most useful for the following applications in a biology lab:

- Measuring bacterial growth in suspension cultures. This is useful, for example, when using engineered bacteria to produce a protein of interest. In that case, the protein production is typically induced when the growth curve reaches the middle of the exponential growth phase (typically at an OD of 0.6).
- Measuring the concentration of particle suspensions, or the turbidity of a liquid sample. This is especially useful for water quality testing, where the turbidity of water is routinely measured.
- Measuring the concentration of solutions of substances that absorb strongly at 590 nm, such a bromophenol blue, pigments/dyes, and chloroplasts.

## 3. Design files

### 3.1 Design Files Summary

All design and code files can be found at https://osf.io/kyu8r/files/ and at https://github.com/czbiohub/ULC-OD-Meter, and are made available under the CERN Open Hardware License, weakly-reciprocal variant, version 2 or later (CERN-OHL-W v2 or later) [1].

The “1-0018 OD Meter” document is a full CAD model of the device, created using the web-based CAD system Onshape (www.onshape.com). Following the link to this document in Table 1 will give full-view access to the model and the ability to export STL files of all the individual parts that need to be 3D printed. Those part files are also available through their respective links in the table. The digits at the start of each file name corresponds to the unique number we have assigned to that part in our internal database of custom parts.

**Table 1:**
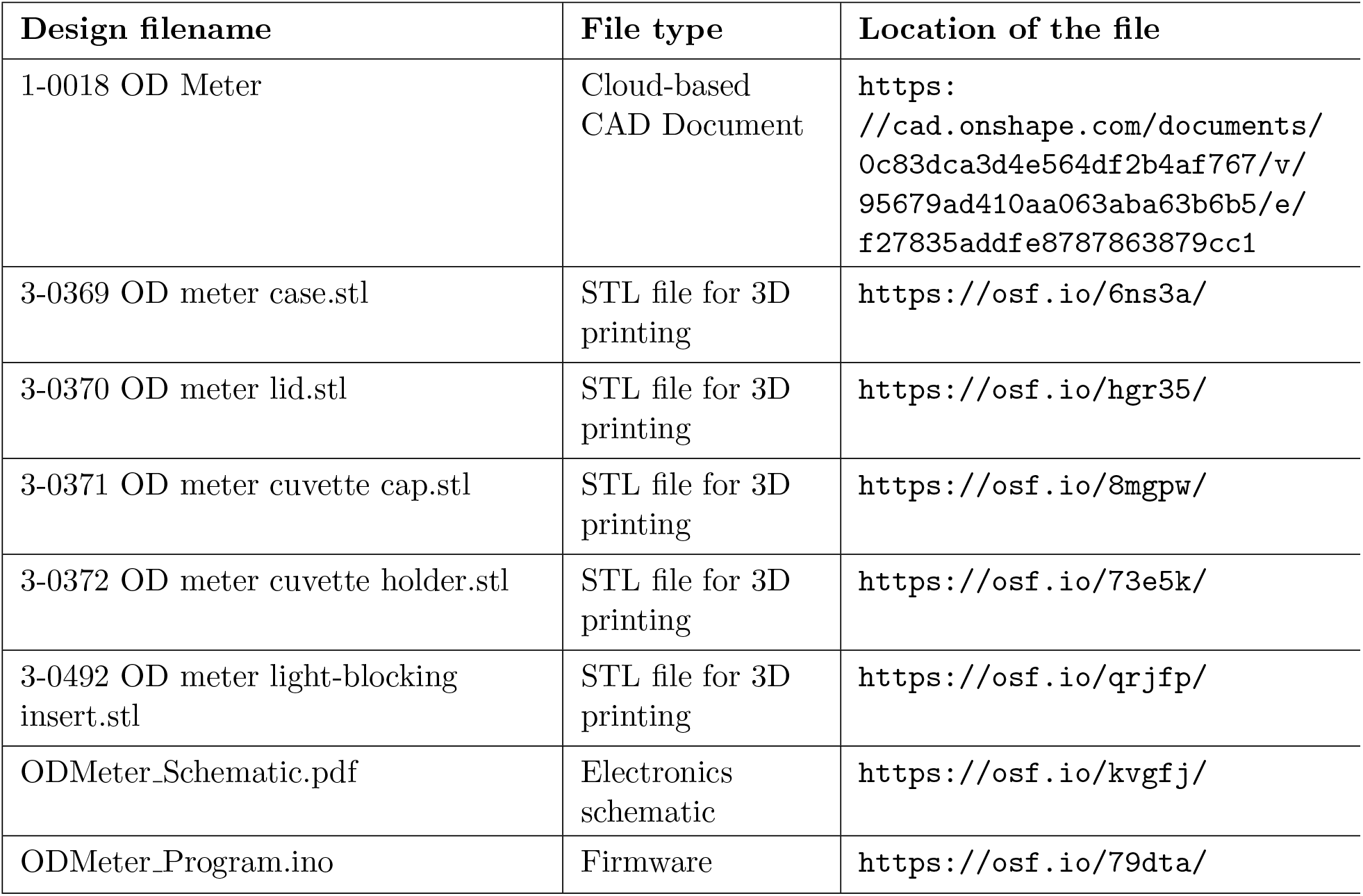
Design files summary.

| Design filename | File type | Location of the file |
| --- | --- | --- |
| 1-0018 OD Meter | Cloud-based CAD Document | https://cad.onshape.com/documents/0c83dca3d4e564df2b4af767/v/95679ad410aa063aba63b6b5/e/f27835addfe8787863879cc1 |
| 3-0369 OD meter case.stl | STL file for 3D printing | https://osf.io/6ns3a/ |
| 3-0370 OD meter lid.stl | STL file for 3D printing | https://osf.io/hgr35/ |
| 3-0371 OD meter cuvette cap.stl | STL file for 3D printing | https://osf.io/8mgpw/ |
| 3-0372 OD meter cuvette holder.stl | STL file for 3D printing | https://osf.io/73e5k/ |
| 3-0492 OD meter light-blocking insert.stl | STL file for 3D printing | https://osf.io/qrjfp/ |
| ODMeter_Schematic.pdf | Electronics schematic | https://osf.io/kvgfj/ |
| ODMeter_Program.ino | Firmware | https://osf.io/79dta/ |

“3-0369 OD meter case.stl” is the file for 3D printing the main body of the instrument case. “3-0370 OD meter lid.stl” is the file for 3D printing the lid for the instrument case.

“3-0371 OD meter cuvette cap.stl” is the file for 3D printing the cap that covers the cuvette during instrument operation, to keep ambient light out of the light detection system.

“3-0372 OD meter cuvette holder.stl” is the file for 3D printing the cuvette holder, which is inserted inside the instrument case during assembly. This holder was build separately from the case to allow easy mounting of the photodiode and LED to the holder.

“3-0492 OD meter light-blocking insert.stl” is the file for 3D printing a cuvette-sized block of black material that is used to calibrate the dark signal of the meter for certain types of samples.

“ODMeter_Schematic.pdf” contains the schematic of the electronics in the device.

“ODMeter_Program.ino” is the Arduino firmware code to run the device. It must be opened and uploaded into the device using the Arduino Integrated Development Environment (IDE).

## 4. Bill of materials

The bill of materials is listed in Table 2. Some of the catalog numbers on the table correspond to packs or boxes containing multiple items (jumper wires, piano wire, screws and nuts). The quantity required (“Qty.” column) is the actual number of individual items used in the build, regardless of the number that come in a pack. The number of items per pack is indicated in parentheses next to the catalog number, while the unit cost on the table is the cost of the pack divided by the number of items it contains. Please note that the cost of the 3D printed components was estimated based on a price of $40 per kilogram of high quality poly-lactic acid (PLA) filament.

**Table 2:** Bill of Materials

| Designator | Catalog # | Qty. | Unit Cost (US\$) | Total Cost (US\$) | Vendor |
| --- | --- | --- | --- | --- | --- |
| 590 nm LED | TLCY5800 | 1 | \$0.61 | \$0.61 | Digi-Key |
| Photodiode | 296-23090-5 | 1 | \$8.80 | \$8.80 | Digi-Key |
| Arduino Micro | 1050-1066 | 1 | \$19.80 | \$19.80 | Digi-Key |
| 16x2 LCD screen | NHD-0216XZ-FSW-GBW | 1 | \$11.63 | \$11.63 | Digi-Key |
| Jumper Wires | 1528-1162 (40/pack) | 22 | \$3.95 | \$3.95 | Digi-Key |
| 1.8 MΩ Gain Resistor | CFR-25JB-52-1M8 | 1 | \$0.10 | \$0.10 | Digi-Key |
| 100 kΩ Gain Potentiometer | 3362S-1-104LF | 1 | \$1.02 | \$0.89 | Digi-Key |
| 100 $\Omega$ LED Resistor | 100EBK | 1 | \$0.10 | \$0.10 | Digi-Key |
| Push Button | 506PB | 1 | \$1.60 | \$1.60 | Digi-Key |
| Header Jumper | S9337 | 1 | \$0.10 | \$0.10 | Digi-Key |
| Mini Breadboard | 1568-1083 | 1 | \$2.95 | \$2.95 | Digi-Key |
| Micro-USB Cable | 380-1431 | 1 | \$2.19 | \$2.19 | Digi-Key |
| 3/64" dia. x 12" long Piano Wire | 8908K52 (100/pack) | 1 | \$0.15 | \$0.15 | McMaster-Carr |
| #2-56 x 3/8" Screw | 92196A079 (100/pack) | 8 | \$0.08 | \$0.64 | McMaster-Carr |
| #2-56 Nut | 91841A003 (100/pack) | 4 | \$0.03 | \$0.12 | McMaster-Carr |
| Enclosure | 3-0369 | 1 | \$3.21 | \$3.21 | 3D printed |
| Enclosure Lid | 3-0370 | 1 | \$1.04 | \$1.04 | 3D printed |
| Cuvette Holder | 3-0372 | 1 | \$0.76 | \$0.76 | 3D printed |
| Cuvette Cap | 3-0371 | 1 | \$0.16 | \$0.16 | 3D printed |
| Light-Blocking Insert | 3-0492 | 1 | \$0.16 | \$0.16 | 3D printed |

### 4.1 Tools needed

The following tools are needed to build the device:

- 3D printer with PLA or other rigid filament
- Soldering iron and lead-free solder
- Hot glue gun with glue sticks
- Tapping wrench with #2-56 tap. Note: If a tap is not available, thread-forming, sheet metal or wood screws of the appropriate diameter (∼0.086” or ∼2 mm) and length can be used to hold the lid, instead of the #2-56 screws listed in the bill of materials.
- Wire cutters
- Cutting blade or wire-stripping tool
- Pliers
- 5/64” hex key
- Optional: Multi-meter for troubleshooting

## 5. Build instructions

### 5.1 Overview

The OD meter presented here consists of a 3D-printed case with a cuvette holder, an Arduino Micro microcontroller, a user interface consisting of a push button and a 16×2 character LCD screen, a 590 nm LED light source, a photodiode with integrated amplifier, and a breadboard for resistors and a potentiometer to set the amplifier gain (See Figure 1). The electronics components are wired together according to the schematic shown in Figure 2.

**Figure 1:**
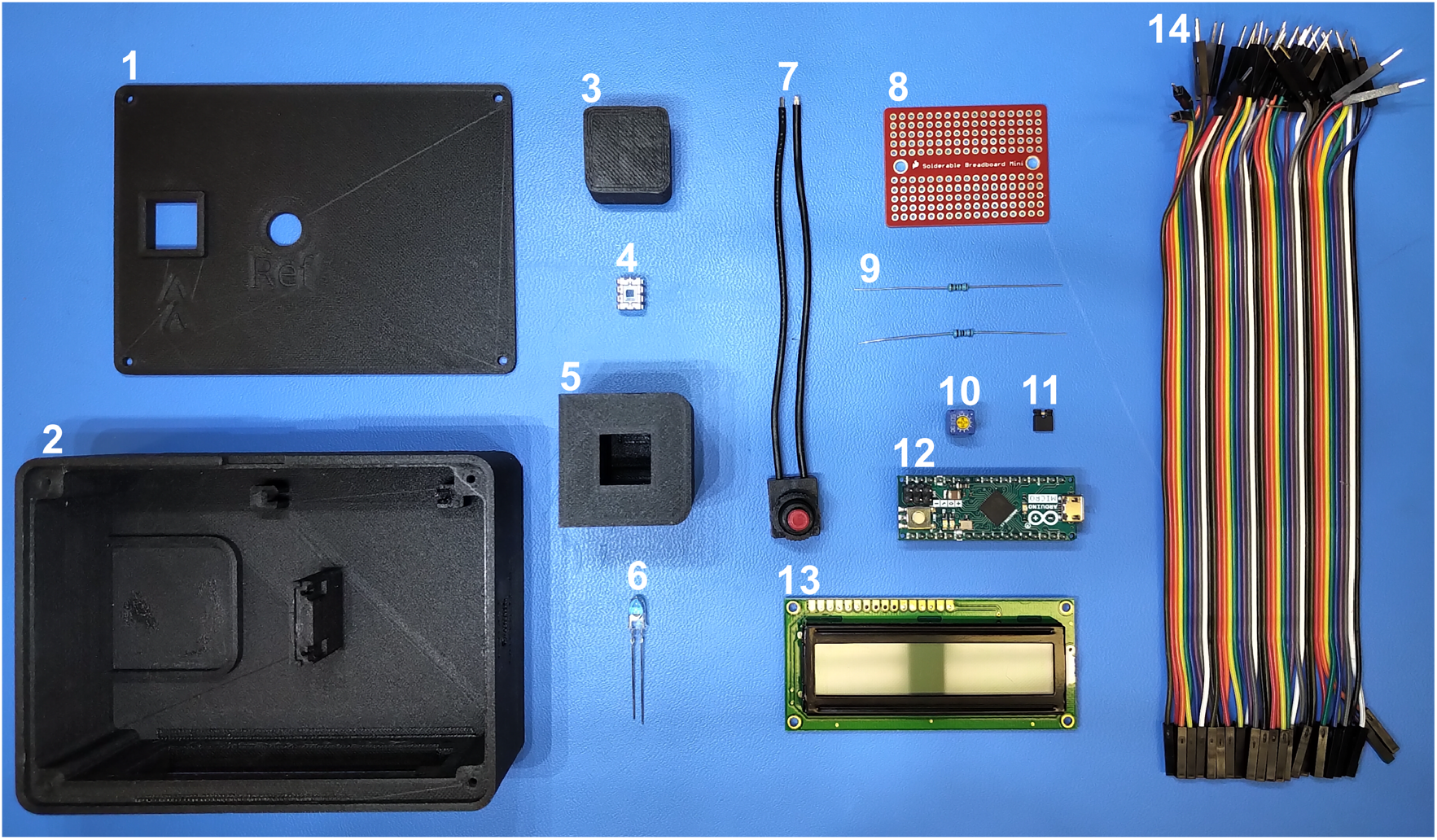
Main components used to assemble the device: (1) 3D printed lid; (2) 3D printed enclosure; (3) 3D printed cuvette cap; (4) photodiode; (5) 3D printed cuvette holder; (6) 590 nm LED; (7) push button; (8) mini breadboard; (9) 1.8 MΩ gain and 100 Ω LED resistors; (10) 100 kΩ gain adjustment potentiometer; (11) header jumper; (12) Arduino Micro; (13) 16×2 LCD screen; (14) jumper wires.

**Figure 2:**
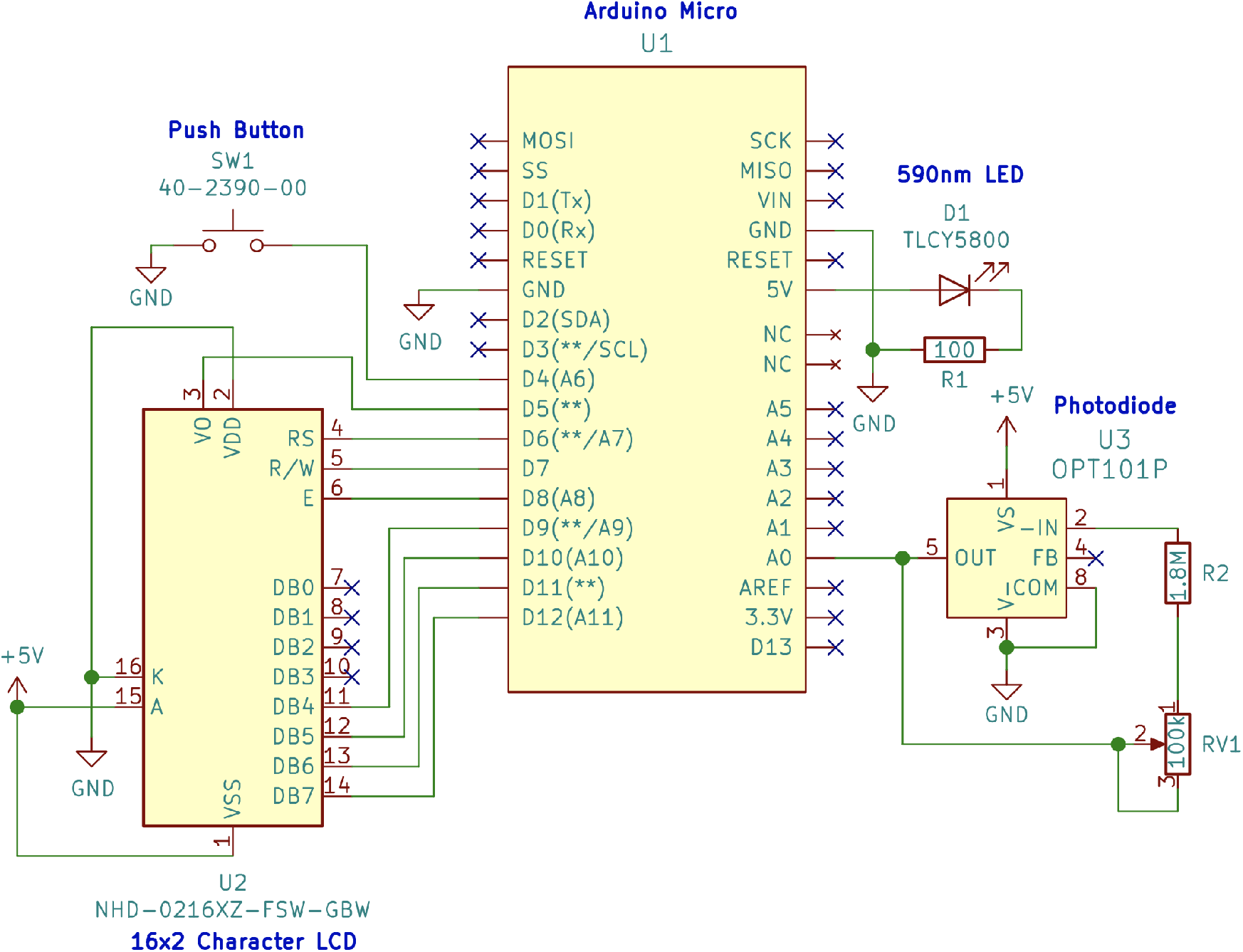
Schematic of the electronics in the device.

### 5.2 Preparation

Prior to assembly, several components need to be prepared. The Enclosure, Enclosure Lid, Cuvette Holder, Cuvette Cap, and Light-Blocking Insert will need to be 3D printed. In addition, wire leads will need to be soldered onto the photodiode and LCD, and the OD meter program will need to be transferred to the Arduino Micro.

#### Enclosure

The Enclosure and related parts (Enclosure Lid, Cuvette Holder, Cuvette Cap, and Light-Blocking Insert) can be printed with a fused deposition modeling (FDM) printer using PLA filament (other rigid printing materials should work as well). We recommend using a matte black material to print the Cuvette Holder and the Light-Blocking Insert to reduce reflections in the measurement unit. If matte black filament is not available, the interior of the Cuvette Holder and the outside of the Light-Blocking Insert can be spray-painted with a black matte paint. Once the parts have been printed, mounting holes for the Arduino Micro in the enclosure should be enlarged with a 1.2 mm diameter (3/64”) drill bit, to receive pins that will hold the Arduino in place. The locations of these holes are highlighted in Figure 3. Insert into each hole a 9.5 mm (3/8”) long steel pin cut from the piano wire, as shown in Figure 3. Finally, threads should be cut into the holes on the four corners of the enclosure using a #2-56 tap (for the screws that will hold the lid). If a tap is not available, thread-forming, sheet metal or wood screws of the appropriate diameter (∼0.086” or ∼2 mm) and length can be used to hold the lid, instead of the #2-56 screws listed in the bill of materials.

**Figure 3:**
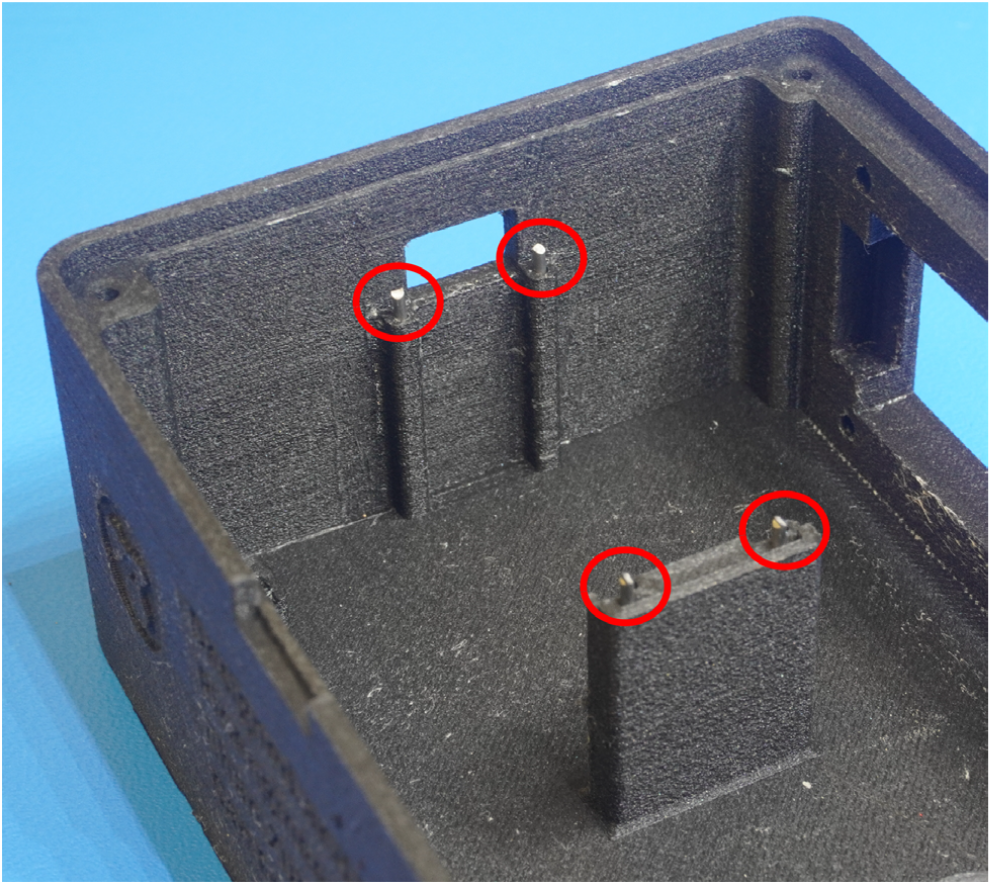
Pins are inserted into four holes in the enclosure to secure the Arduino Micro to the it.

#### Soldering wire leads onto components

Separate nine jumper wires from the bundle and cut off the female connectors, leaving only a male connector on each wire. Separate eight jumper wires from the bundle and cut off the male connector, leaving only a female connector on each wire. Strip off 3 mm of insulation from the cut end of all the wires and deposit a small amount of solder onto all of the stripped ends with a soldering iron. Deposit a small amount of solder onto pins 1, 2, 3, 5, and 8 of the photodiode. IMPORTANT: Do not leave the soldering iron in contact with the pins for too long, as the photodiode can be damaged by the heat. Connect a wire with male connector onto each one of the photodiode pins that received solder. Follow an identical procedure with pins 1-6 and 11-16 on the LCD. LCD pins 1, 2, 15, and 16 receive wires with male connectors, while pins 3-6 and 11-14 receive wires with female connectors. See Figure 4 for the end result of the soldering.

**Figure 4:**
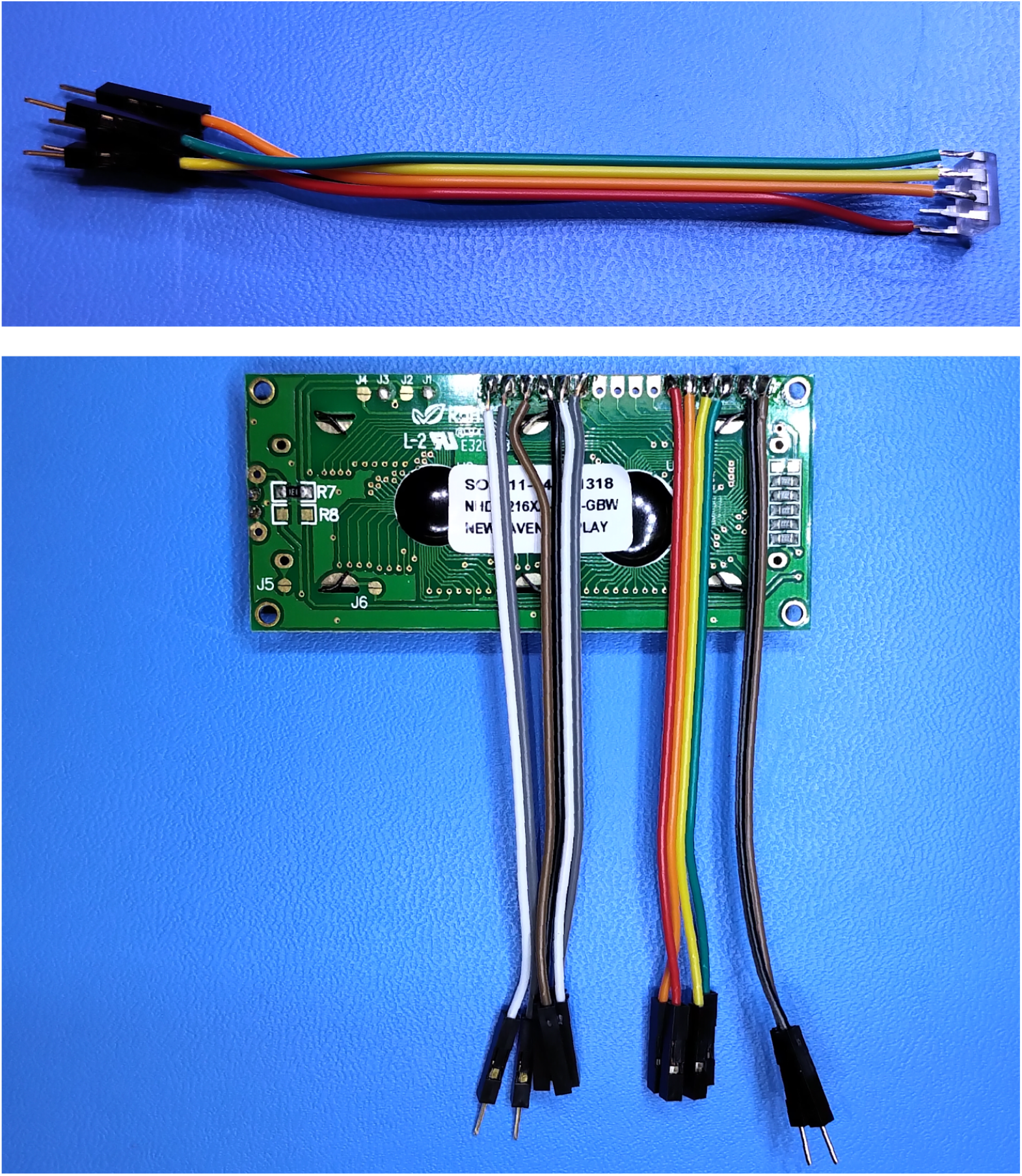
Jumper wires are soldered directly to the pins of the photodiode (top) and LCD (bottom). All wires soldered to the photodiode have male connectors on their free end. LCD pins 1, 2, 15, and 16 receive wires with male connectors on their free end, while pins 3-6 and 11-14 receive wires with female connectors.

#### Arduino Micro programming

The OD meter program (file “ODMeter Program.ino”) must be transferred to the Arduino Micro using the Arduino IDE. Version 1.8.9 of the IDE was used here, but other versions may work as well. Once the IDE has been downloaded, the LiquidCrystal library will need to be installed into it. To install the library select the “Sketch” drop-down menu and select “Manage Libraries…” from the “Include Library” section. From here, search for the LiquidCrystal library and install version 1.0.6. Connect the Arduino to the computer with a micro-USB cable. In the “Tools” drop-down menu, set the board option to “Arduino/Genuino Micro” and set the port option to the serial port hosting the Arduino. Open the program file, “ODMeter Program.ino”, and click the “Upload” button to transfer the program to the Arduino. To monitor the output of the system from a computer rather than the LCD on the system, the Arduino IDE includes a serial monitor function. This will be used when aligning the photodiode in the measurement module and can also be used to help troubleshoot display issues, if encountered.

### 5.3 Breadboard

The breadboard has two banks of thru-holes (top and bottom), each with 17 columns and 5 rows of holes. The top and bottom banks of holes are isolated from each other. Holes in a column are electrically connected to each other if they are in the same bank (top or bottom), while columns are isolated from each other. Follow Figure 5 to populate the breadboard with components and establish three solder bridges, ensuring that connections follow the schematic in Figure 2. Make a solder bridge across two columns to create the 5 V rail (red rectangle in the figure), a solder bridge across two other columns to create the GND rail (black rectangle), and a solder bridge connecting the two left-most pins of the potentiometer. Finally, solder the male connectors of six jumper wires to the breadboard as follows: Two jumper wires to the 5 V column, one wire to the GND column, one wire to the end of the 100 Ω LED resistor (column L, orange), one wire to the middle pin of the potentiometer (column O, blue), and one wire to the B column (yellow). Solder one wire of the push button to the B column (yellow), and the other wire to the GND rail. Using the schematic on Figure 2 as a guide, connect the jumper wire on the GND column to a GND pin on the Arduino, connect one jumper wire on the 5 V column to the 5 V pin on the Arduino, connect the jumper wire attached to the middle potentiometer pin (O, blue) to the A0 pin on the Arduino, and connect the jumper wire on the B column (yellow) to pin D4 on the Arduino. Short the AREF and 3V3 pins on the Arduino with the header jumper to set the reference analog voltage to 3.3 V.

**Figure 5:**
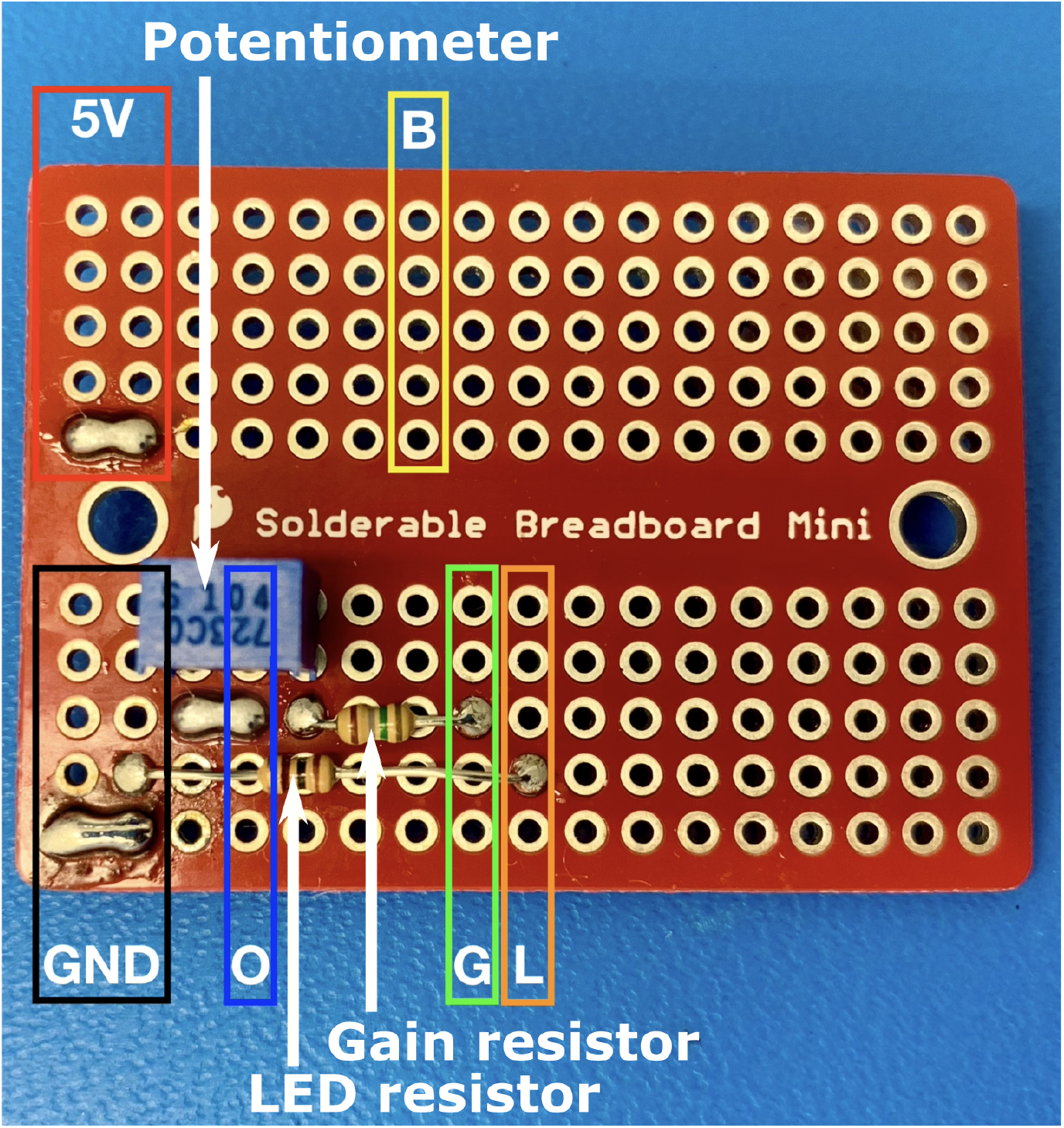
Components soldered to the breadboard. The 5 V rail (red rectangle) is created with a solder bridge across two columns on the top bank of holes, while the GND rail (black) is extended with a solder bridge across two columns on the bottom bank. A solder bridge connects the two left-most pins of the potentiometer. Column O (blue) will be connected to pin 5 of the photodiode. Column G (green) will be connected to pin 2 of the photodiode. Column L (orange) will be connected to the cathode of the LED. Column B (yellow) will be connected to one wire of the push button and to pin D4 on the Arduino Micro.

### 5.4 Light source and light sensor

Using Figure 5 as a guide, solder the photodiode wires and the LED to the breadboard as follows: Photodiode pins 3 and 8 to the GND rail, pin 1 to the 5 V rail, pin 2 to column G (green), and pin 5 to column O (blue), LED anode (positive terminal) to the unused jumper wire on the 5 V rail, and LED cathode (negative terminal) to the jumper wire soldered to column L (orange).

Connect the Arduino to a USB port on a computer to power the device and confirm correct wiring of the power rails and the polarity of the LED. If the LED does not turn on, disconnect the Arduino from the USB port and double-check all the wiring to make sure it conforms exactly to the schematic in Figure 2. If the LED turns on, wait 20 min for it and the photodiode to stabilise, then proceed to mount them as shown in Figure 6. Proper alignment of the LED and photodiode is critical for the performance of the device. Improper alignment will reduce the signal-to-noise ratio and dynamic range of the OD measurements. Place the photodiode against its rectangular opening on the side of the Cuvette Holder. Dispense hot glue around the photodiode and maintain pressure on the photodiode as the glue cools to preserve alignment. Insert the illuminated LED into the proper aperture of the Cuvette Holder. Center the beam projected by the LED onto the photodiode aperture and fasten the LED with hot glue. A thin white piece of paper can be inserted into the Cuvette Holder opposite to the LED to visually confirm maximal brightness projection onto the photodiode. Maintain pressure on the LED until the glue has solidified to preserve alignment. Launch the serial monitor in the Arduino IDE to read the output of the photodiode. Adjust the potentiometer on the breadboard until the output reads approximately 1015 counts in ambient light conditions. The output may need about 45 minutes to settle after adjusting the potentiometer. Fix the potentiometer knob with hot glue to prevent it from drifting over time.

**Figure 6:**
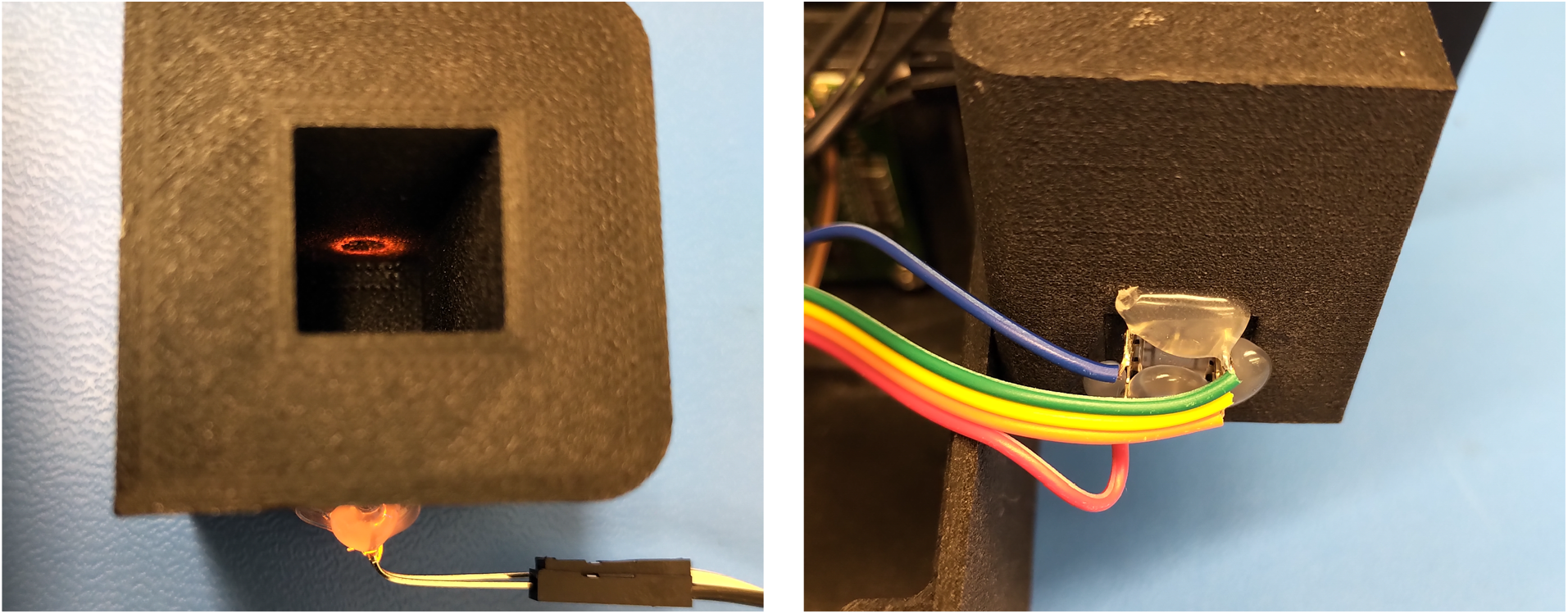
The photodiode is first aligned such that the emitted light is centered over the photodiode aperture. Once aligned, the component is secured with hot glue and is held in place until the glue solidifies. Afterwards, the position of the LED is adjusted until maximum signal is obtained from the photodiode. The LED is then secured with hot glue and is held in place until the glue solidifies.

### 5.5 Screen

Disconnect the Arduino from the computer and solder the LCD to the breadboard. Pins 1 and 15 are soldered to the 5 V rail on the breadboard, while pins 2 and 16 are soldered to the GND rail. Next, connect LCD pins 3-6 and 11-14 to Arduino pins D5 to D12, in consecutive (ascending) order. Reconnect the Arduino to the computer to confirm that the display is functioning - the LCD should display the same values that are received through the serial port on the Arduino IDE. If the LCD contains obscure characters or if the characters are not visible, confirm that the wiring connections conform to the schematic in Figure 2. If the connections are correct, a wire may have been damaged and is not connected internally, or a solder joint might be defective. Use a digital multi-meter and probe for discontinuities in the connections, if needed. Confirm that the button functions correctly. When the button is pressed the screen should display messages according to the operation instructions. If everything is working properly, disconnect the Arduino from the computer to turn the device off and move on to the packaging section.

### 5.6 Packaging

Once all components have proven to be functional, they can be mounted into the enclosure, as shown in Figure 7. Start by fastening the LCD to the enclosure with #2-56 socket head screws and nuts. Next, insert the Cuvette Holder into its cradle inside the enclosure. Afterwards, insert the breadboard into its holding slots and secure it with a drop of hot glue. Finally, align the holes in the Arduino board with the steel dowel pins and firmly press onto the support structures below. Fix the Arduino by dispensing dots of hot glue along the support structures. All the places inside the enclosure that require hot glue are indicated in Figure 8. Put the lid onto the enclosure and fasten it with #2-56 screws that are tightened very lightly. Over-tightening the lid screws will damage the threads cut into the enclosure.

**Figure 7:**
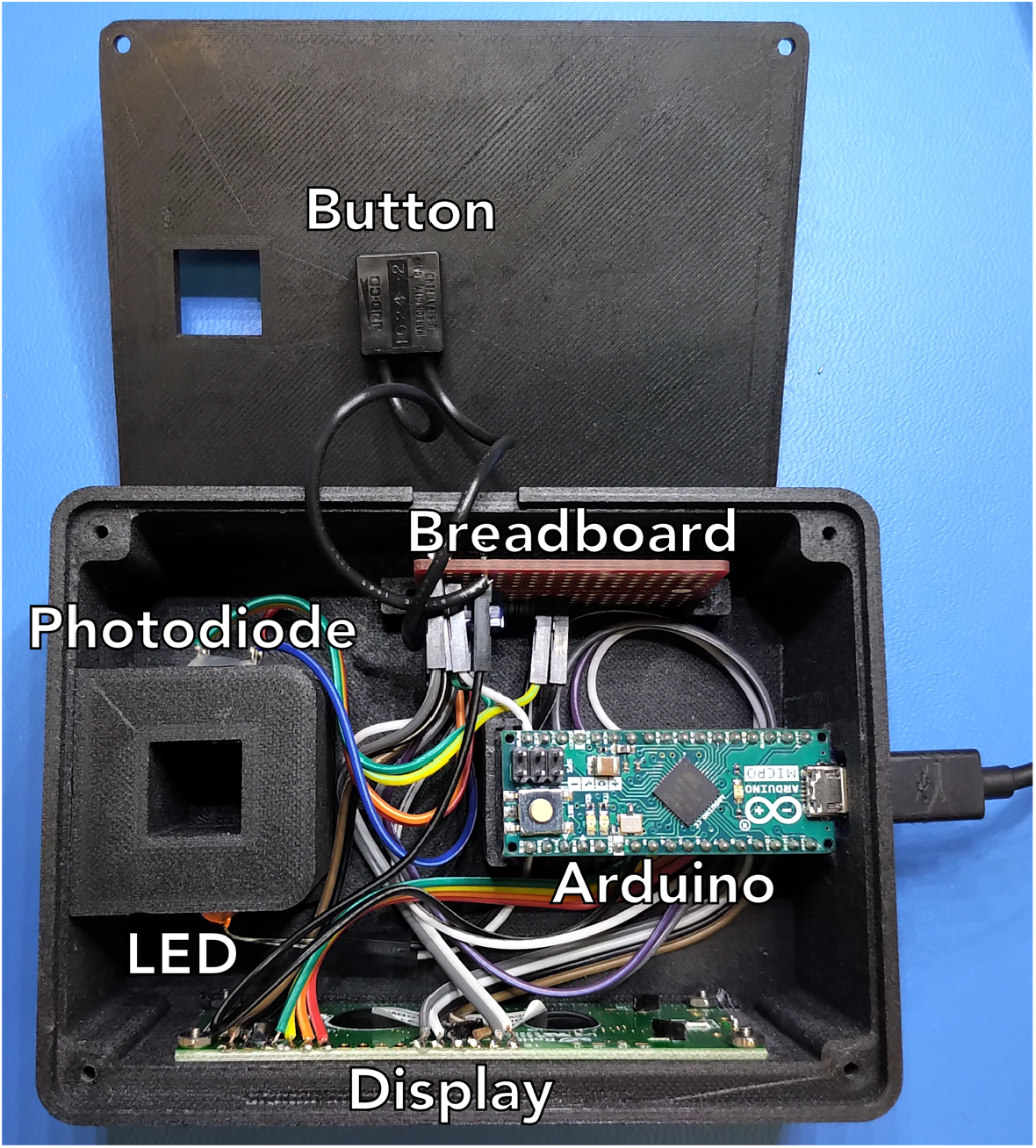
Location of all major components inside the enclosure.

**Figure 8:**
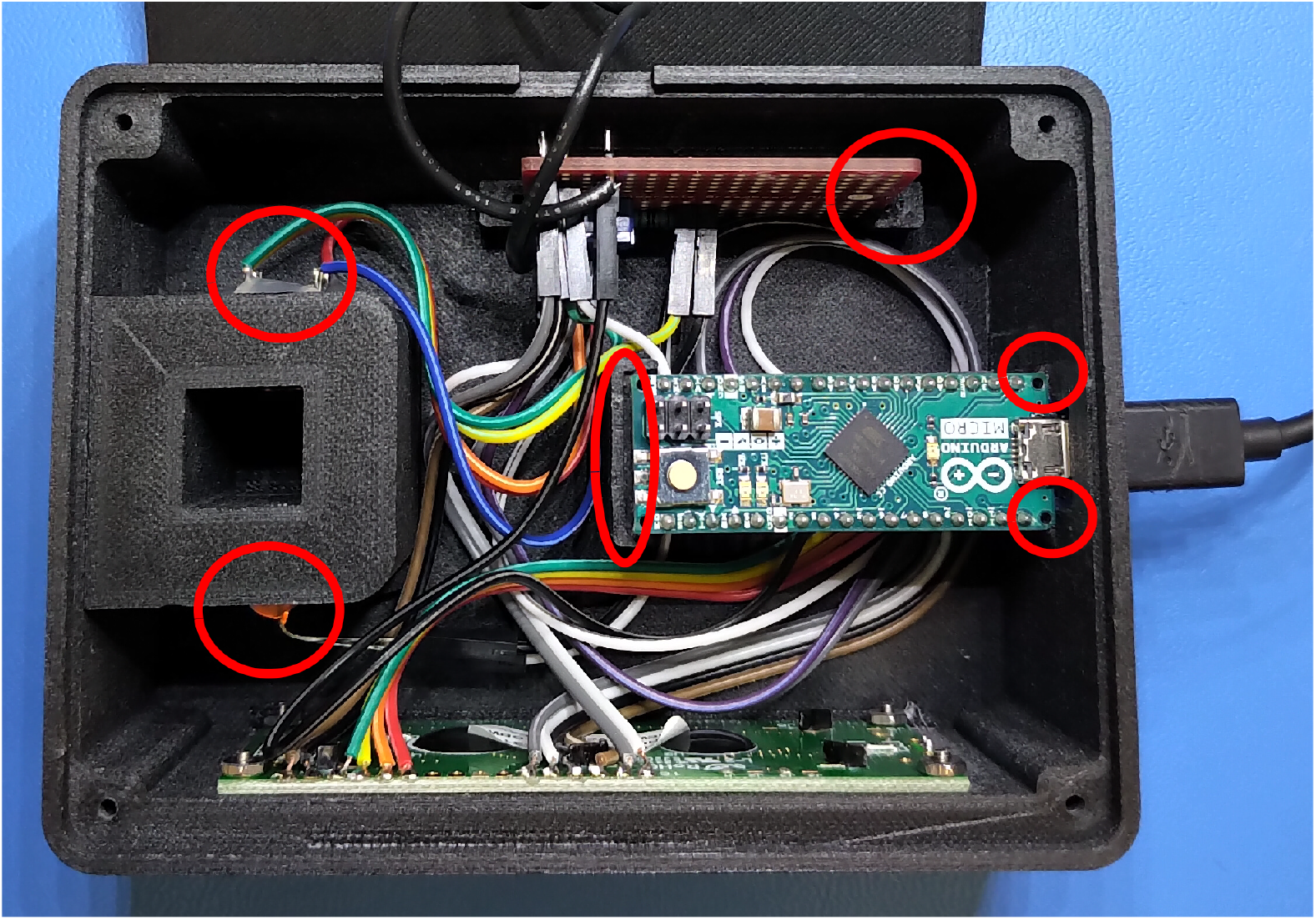
Highlighted areas indicate where hot glue will be required to fasten components to the enclosure and to the cuvette holder.

## 6. Operation instructions

The device has two modes of operation: (A) “Raw” mode, where where the screen only displays a number between 0 and 1024, proportional to the raw light intensity measured by the photodiode and (B) “OD” mode, where both OD and raw light intensity values are displayed. Optical density is defined by eq. 1, where *I*_*blank*_ is the light intensity measured with a “blank” cuvette (filled with the solvent/medium that will contain the bacteria or light absorbing substance to be measured), *I*_*dark*_ is the light intensity measured with the Light Blocking Insert, or a cuvette filed with a very high OD sample of the same type being measured, depending on the situation, and *I*_*samp*_ is the light intensity measured with the sample cuvette.

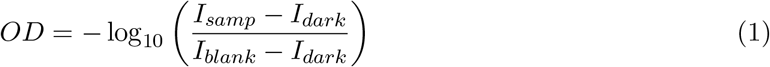

To operate the device, connect it to a powered USB port on a computer, USB charger or battery pack and let the machine warm up for at least 20 minutes before performing any measurements. The device always starts in “Raw” mode when it is first powered up. Expect to see the raw intensity values fall and gradually stabilize as the device warms up. The operation mode is switched by pushing the “Ref” button on the lid, according to the following sequence:

1. In “Raw mode” press the “Ref” button to go into “OD” mode.
2. Device asks for the dark reference.
3. Insert the dark reference cuvette or Light-Blocking Insert into the device and cover it with the cap.
4. Press “Ref”.
5. Device records *I*_*dark*_ and saves it to memory, then asks for the blank cuvette.
6. Insert the blank reference cuvette into the device and cover it with the cap.
7. Press “Ref”.
8. Device records *I*_*blank*_ and saves it to memory.
9. Device enters “OD mode” and starts displaying both OD and raw intensity values, as shown in Figure 10.
10. In “OD mode”, press “Ref” for at least 1 second to re-measure the dark and blank references with the same sequence as above.

Note that the *I*_*blank*_ and *I*_*dark*_ values are lost when the device is powered off. Always be mindful of the light direction (indicated by the symbol ≫ on the lid) when inserting the cuvettes. You have to decide in advance if you want the light traveling through the thick or thin orientation of the cuvettes, depending on how opaque the fluid being measured is. The sample cuvettes should be inserted in exactly the same orientation relative to the light direction as the reference and dark cuvettes. Make sure to always cover the cuvette with the cap when measuring to keep ambient light from affecting the measurement.

**Figure 9:**
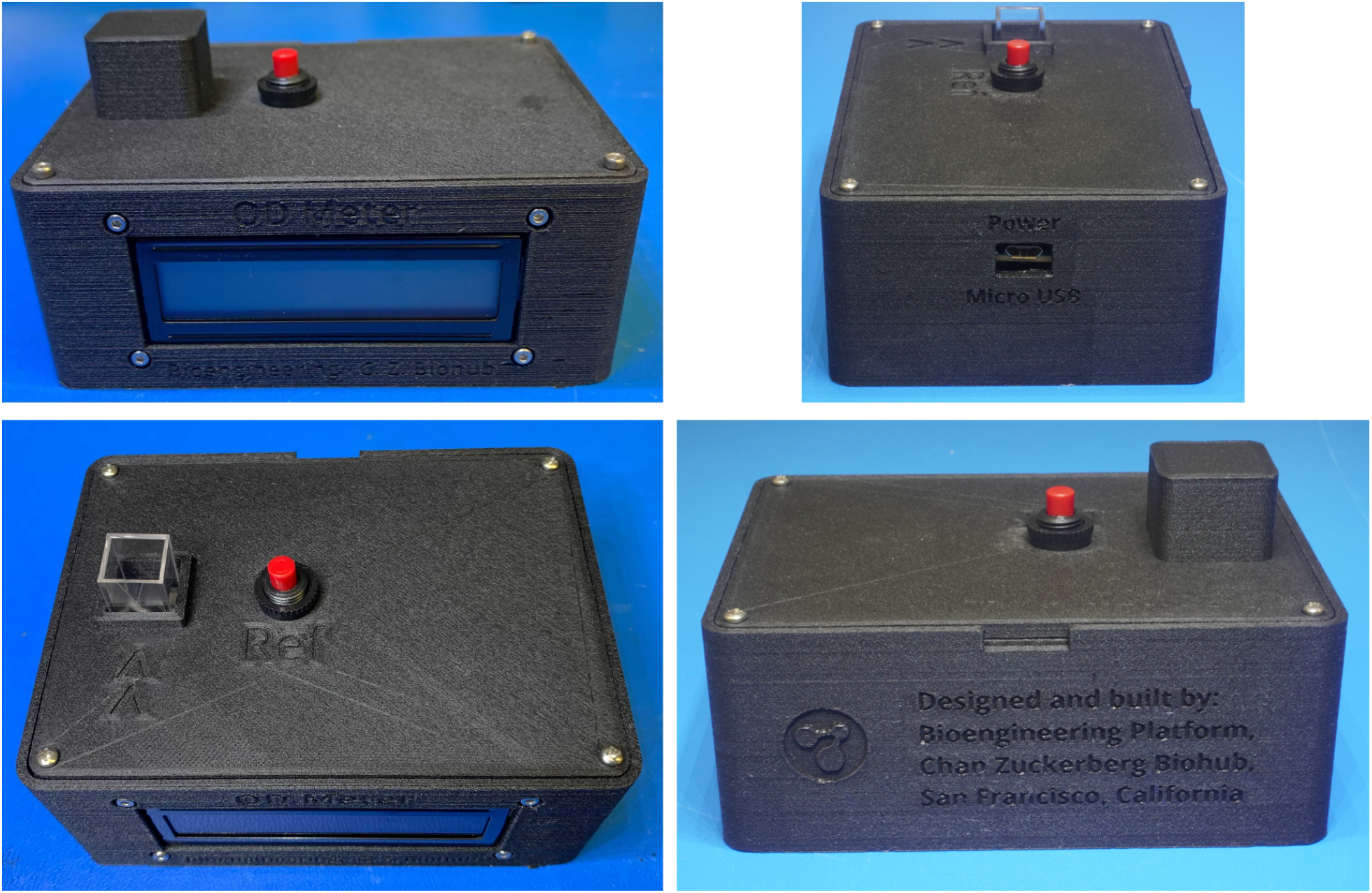
Fully assembled device.

**Figure 10:**
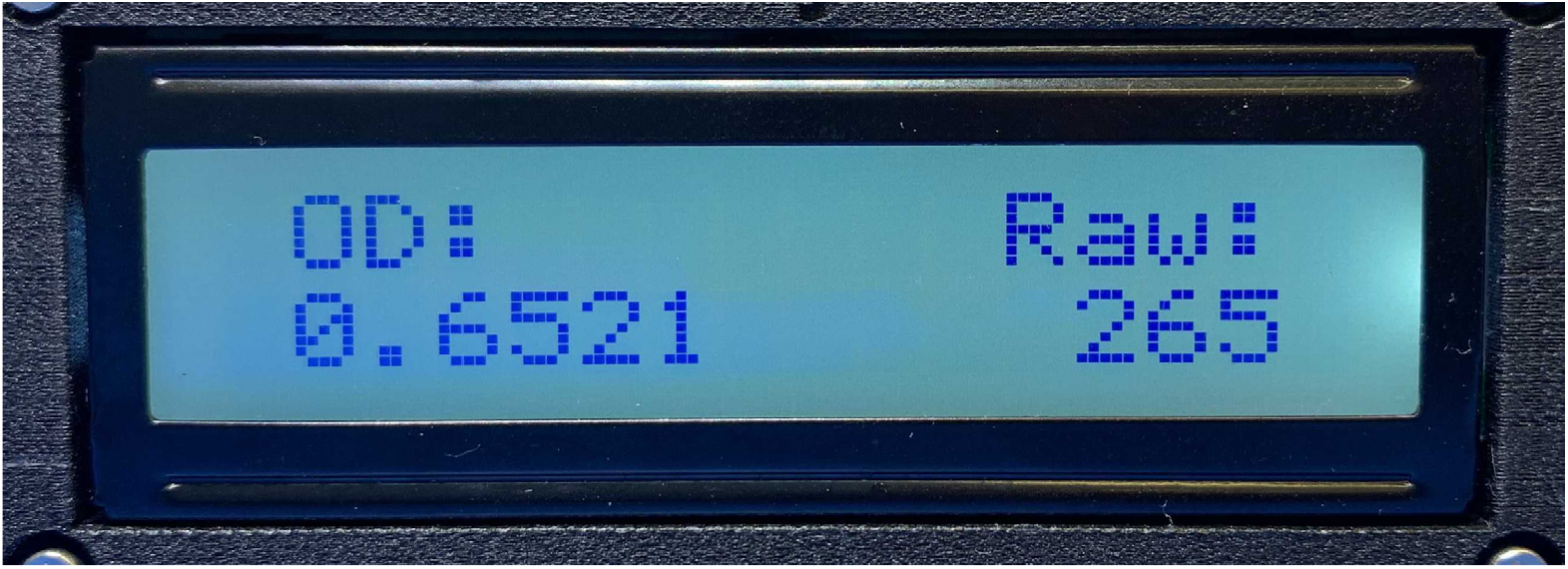
Device screen in OD mode, showing OD and raw intensity values.

For measurements of purely absorbing (non-scattering) solutions up to OD 1.0 and bacterial suspensions (scattering) up to OD 2.0, the dark reference can be easily measured with the 3D-printed light-blocking insert (part# 3-0492). However, in order to measure OD values above 1.0 for purely absorbing (non-scattering) solutions such as dye, the dark reference measurement is more nuanced. Using the Light-Blocking Insert as a dark reference for dye gives a dark value that is too low and will result in underestimation of OD values above 1.0. When attempting to measure solutions with OD values above 1.0 the dark reference must be acquired with a cuvette filled with the same type of solution being measured, at a concentration such that the OD is well above the range being measured, but not as high as the OD of the Light-Blocking Insert. Our observations with purely absorbing samples suggest that a small amount of light reaches the sensor independently of dye concentration, presumably caused by off-axis light rays coupling through the cuvette walls by total internal reflection (TIR), resulting in a increased dark signal. Notably, this effect is not observed with highly scattering samples such as bacterial suspensions, consistent with the hypothesis that TIR may be frustrated in the presence of higher refractive index components in solution, leading to a more complete extinction of the light. Such off-axis light is present in the system because the device does not have any optical elements to collimate the light from the LED light source, and relies instead on narrow LED and sensor pupils to reduce off-axis light. The above limitations at high OD are compounded by the 10-bit ADC in the Arduino, which places a hard cap on the dynamic range.

### 6.1 Dark reference calibration

The procedure for calibrating the dark reference for precise OD measurements of solutions above 1.0 is as follows:

1. Prepare a solution of the substance of interest at a concentration that gives a raw reading of approximately 70 in the OD meter. This reading is the first guess for *I*_*dark*_.
2. Perform a dilution series of the same solution prepared in step 1, with each step diluting by 10 to 20%. Measure the raw light intensity of each dilution step. A minimum of 10 dilution steps is recommended.
3. Measure the raw light intensity of solvent alone in the cuvette. This reading is *I*_*blank*_.
4. Convert all the measured raw values to OD using eq. 1 and plot them as a function of the dilution level.
5. If the plot is a straight line, the *I*_*dark*_ value measured in step 1 is the optimal one. The straightness can be evaluated using a linear fit. For future experiments using this same substance, the dark reference can be acquired with the same concentration of the substance used in step 1.
6. If the plot is not a straight line, recalculate the OD values with a range of values of *I*_*dark*_ to find the value of *I*_*dark*_ that makes the plot as linear as possible (performing a linear regression fit to the data points is the best way to assess linearity). For future experiments using this same substance, the dark reference can be acquired with a concentration of the substance that gives this optimal *I*_*dark*_ raw value.

## 7. Validation and characterization

The initial characterization of the device was done by measuring the OD of three dilution series of blue food dye in water (labeled D1, D2, and D3) with both the OD meter and a SpectraMax M3 spectrophotometer set to 590 nm (Molecular Devices, LLC, San Jose, California, USA). At each step in the dilution series, the dye concentration was reduced by 20%, down to OD values of 0.08, 0.07, and 0.05 for D1, D2, D3, respectively. Figure 11 shows three dilution curves of blue food dye, with the SpectraMax OD value in the X axis and the low-cost device OD value in the Y axis. For two of the curves (samples D1 and D2), the dark reference was measured with cuvettes filled with a concentration of blue dye that gave a raw light intensity value of 70 for sample D1 and 73 for sample D2. For sample D3 the dark reference was acquired with the Light-Blocking Insert, giving a raw light intensity reading of 22, which results in the OD being underestimated for values above ∼1.1 (Figure 11A), for the reasons explained in the “Operation instructions” section. For that sample, the optimal dark reference value is 60, obtained by following steps 4-6 in the dark reference calibration procedure. Figure 11B illustrates how collecting the dark reference with the 3D printed light-blocking insert is good enough when measuring OD values up to 1.1. In this range, the underestimation of OD in sample D3 due to the incorrect *I*_*dark*_ calibration is not significant.

**Figure 11:**
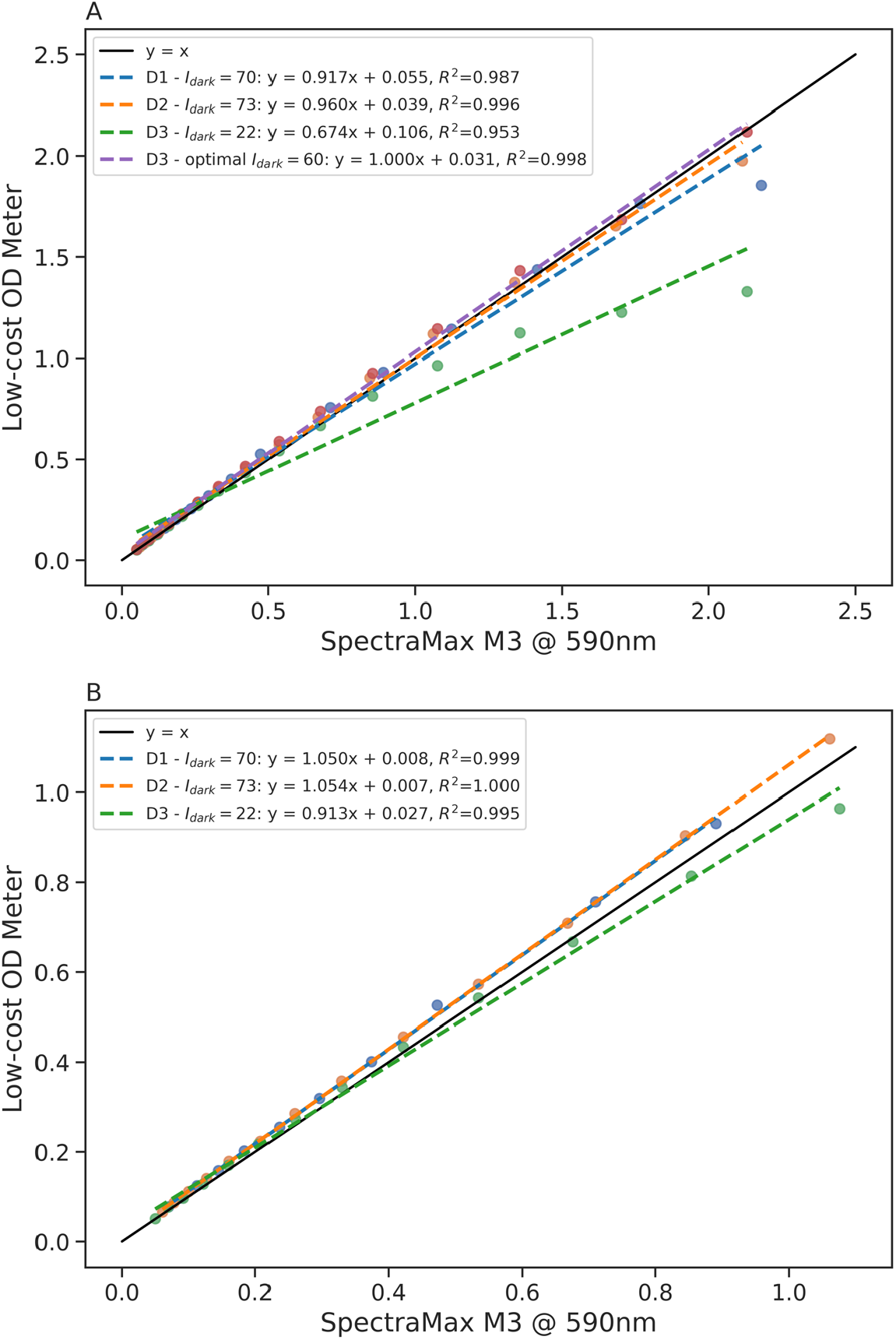
Blue food dye dilution series for three samples (D1, D2, and D3), comparing OD values obtained from the OD meter to values from a SpectraMax M3 spectrophotometer at 590 nm. At each step in the dilution series, the dye concentration was reduced by 20%, down to OD values of 0.08, 0.07, and 0.05 for D1, D2, D3, respectively. The dark references for samples D1 and D2 were acquired with cuvettes filled with blue dye (*I*_*dark*_ = 70 and 73, respectively). The dark reference for sample D3 was acquired using the 3D printed light blocking insert (*I*_*dark*_ = 22). The blank reference for all samples was acquired with pure water in a cuvette (*I*_*blank*_ = 1000, 1019, 980, for D1, D2, D3, respectively). Dashed lines are linear fits to the data points of the same color up to the OD range of each plot. (A) OD values up to 2.2. In this range the *I*_*dark*_ calibration for D3 leads to underestimating the OD values above OD 1.1 (green points and line). The optimal *I*_*dark*_ for this sample is 60, which properly linearizes the curve (purple points and line). (B) OD values up to 1.1. In this range the underestimation of OD in sample D3 due to the incorrect *I*_*dark*_ calibration is not significant.

Figure 12 shows plots of the OD measured with the low cost device on two suspensions of *E. coli* in Luria-Bertani (LB) broth that started at different concentrations (the bacteria were cultured at 37°C). The concentration was reduced by 20% at each step in the series by diluting with fresh LB broth. For both samples the 3D printed light blocking insert was used as a dark reference, and the blank reference was acquired with pure LB broth in a cuvette. It is very clear from these plots that the OD meter can properly measure OD values of bacterial suspensions from 0.05 to 2.0 by just using the 3D printed light blocking insert as a dark reference.

**Figure 12:**
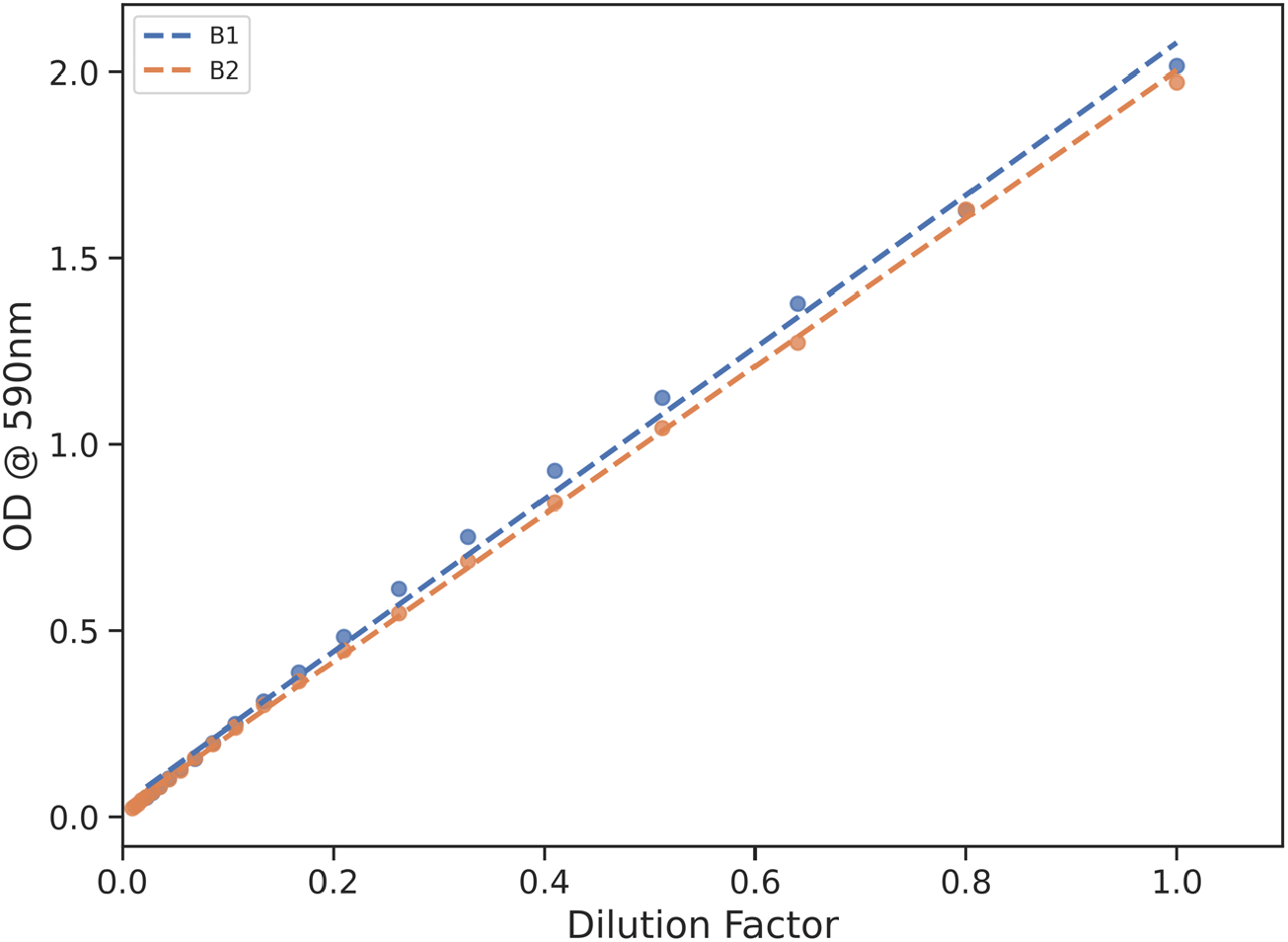
*E. coli* dilution series for two samples (B1 and B2, in LB broth), plotting OD values obtained from the OD meter as a function of the dilution factor. Each sample was started at a slightly different bacterial concentration (OD 2.02 for B1, 1.97 for B2), and the bacterial concentration was reduced by 20% at each step in the series by diluting in fresh LB broth, down to an OD value of 0.05 for B1, and 0.02 for B2. The dark references for both samples were acquired with the 3D printed light blocking insert (*I*_*dark*_ = 28 for B1, 31 for B2), and the blank reference was acquired with pure LB broth in a cuvette (*I*_*blank*_ = 962 for B1, 971 for B2). Dashed lines are linear fits to the data points.

To summarize:

- The device can accurately measure OD values of solutions from about 0.05 up to 1.0 using the 3D printed light blocking insert as a dark reference.
- Measuring OD values of solutions between 1.0 and 2.0 requires a calibration process to determine the correct dark reference.
- The device can accurately measure OD values of bacterial suspensions up to 2.0 using the 3D printed light blocking insert as a dark reference.

## 8. Acknowledgements

The authors would like to thank Vida Ahyong for asking us to create a simple and cheap OD meter to help with her CRISPR training classes in Uganda, and to Chaz Langelier for carrying the device in his luggage all the way to Kampala. This work was supported by the Chan Zuckerberg Biohub.

## 10. Human and animal rights

This work did not involve any human or animal experimental subjects.

